# Protein language models learn evolutionary statistics of interacting sequence motifs

**DOI:** 10.1101/2024.01.30.577970

**Authors:** Zhidian Zhang, Hannah K. Wayment-Steele, Garyk Brixi, Haobo Wang, Matteo Dal Peraro, Dorothee Kern, Sergey Ovchinnikov

## Abstract

Protein language models (pLMs) have emerged as potent tools for predicting and designing protein structure and function, and the degree to which these models fundamentally understand the inherent biophysics of protein structure stands as an open question. Motivated by a discovery that pLM-based structure predictors erroneously predict nonphysical structures for protein isoforms, we investigated the nature of sequence context needed for contact predictions in the pLM ESM-2. We demonstrate by use of a “categorical Jacobian” calculation that ESM-2 stores statistics of coevolving residues, analogously to simpler modelling approaches like Markov Random Fields and Multivariate Gaussian models. We further investigated how ESM-2 “stores” information needed to predict contacts by comparing sequence masking strategies, and found that providing local windows of sequence information allowed ESM-2 to best recover predicted contacts. This suggests that pLMs predict contacts by storing motifs of pairwise contacts. Our investigation highlights the limitations of current pLMs and underscores the importance of understanding the underlying mechanisms of these models.

**Significance Statement:** Protein language models (pLMs) have exhibited remarkable capabilities in protein structure prediction and design. However, the extent to which they comprehend the intrinsic biophysics of protein structures remains uncertain. We present a suite of analyses that dissect how the flagship pLM ESM-2 predicts structure. Motivated by a consistent error of protein isoforms predicted as structured fragments, we developed a completely unsupervised method to uniformly evaluate any protein language model that allows for us to compare coevolutionary statistics to older linear models. We further identified t hat E SM-2 a ppears to have a precise context size that is needed to predict inter-residue contacts. Our study highlights the current limitations of pLMs and contributes to a deeper understanding of their underlying mechanisms, paving the way for more reliable protein structure predictions.

---

**D**etermining the structure of a protein is a critical first step to understanding its function in biology; therefore, tremendous efforts have been devoted to the task of predicting protein structure from sequence. AlphaFold2 (AF2) (1) dramatically improved the prediction accuracy of single protein structures in the CASP14 challenge. Central to AF2’s methodology are multiple sequence alignments (MSA) that contain information on evolutionary couplings between amino acids within a structure. However, proteins folding in solution know nothing of their evolutionarily-related counterparts, and methods that can accurately predict structure from a single sequence alone would ideally bring us closer to understanding the biophysics of protein folding. Furthermore, using MSAs to predict structure limits the usefulness of these methods in contexts where few sequence homologs are available. These motivations have driven the development of single-sequence, i.e. MSA-free, structure prediction methods, such as OmegaFold (2), RGN2 (3), and ESMFold (4). Many of these methods fundamentally differ in their training from AF2 and RoseTTAFold (5): rather than being trained in a supervised manner to predict structure, they were developed initially from models trained using the unsupervised task of masked language modelling. OmegaFold is based on the protein language model OmegaPLM, RGN2 is based on the language model aminoBERT, and ESMFold is based on the protein language model ESM-2. Given that these methods do not require MSAs as input, this has raised the question, have protein language models learned the intrinsic physics of folding a single amino acid sequence? More generally, how do they achieve high predictive accuracy from a single sequence? A deeper understanding and interpretation of these models is needed for them to be used reliably.

In this work, we dissected how the language model ESM-2 enables highly accurate structure prediction by evaluating three different hypotheses for its function (Fig. 1). We start with the hypothesis that ESM-2 truly has learned protein folding from physics. This is already contradicted by the result that ESM-2 performance is highly correlated with the number of sequence neighbors in the training set across all model sizes (4). If ESM-2 truly had learned the physics of protein folding, its performance should not depend on the number of sequence neighbors of a given protein. This hypothesis was further contradicted by a striking consistent error we observed in structure predictions of protein isoform sequences that suggested ESM-2 was erroneously using coevolutionary information to predict isoform sequences in the full-length context of a protein. Based on this finding, we formulated two alternate hypotheses: ESM-2 is performing a “full fold lookup”, it has memorized a collection of folds, and given an entire protein sequence it matches contact predictions to a particular fold class. A third hypothesis is that it has memorized motifs represented as pairs of fragments. We designed a series of experiments to test these two hypotheses, and provide evidence for the third hypothesis: that ESM-2 has learned pairwise dependencies conditioned on sequence motifs and the relative separation between the sequences.

**Fig. 1.**
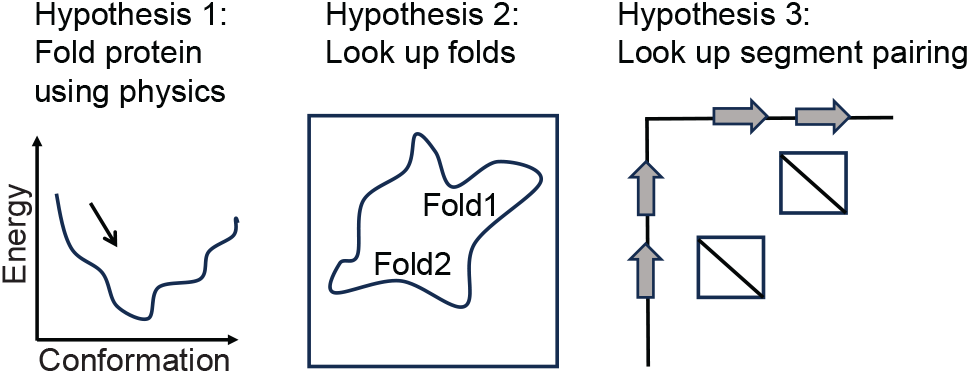
Three hypotheses of how language models predict protein structures.

## Results

### Language models predict unrealistic structures for protein isoforms

Protein isoforms resulting from splicing events within structured domains have long been presented as a pathology for homology-based structure modelling (6–9), since their sequences are very similar to their full-length proteins, yet are often likely unfolded and nonfunctional(9). These isoforms offered an opportunity to evaluate the capabilities of the current protein structure prediction methods. If state-of-the-art protein structure prediction approaches predict such isoforms as either unfolded or alternately structured, it would imply an intrinsic understanding of the biophysics of protein folding. We curated a dataset of 18 domain-splitting isoforms that had previously been identified in references (6–9), and made structure predictions using AlphaFold2 (with MSA input), OmegaFold (language model), and ESMFold (language model) (see Methods).

An example isoform from human myoglobin, first discussed as an example of this phenomenon in (8), is depicted in Fig. 2A. The isoform’s predicted structures in AF2, OmegaFold, and ESMFold have 0.49, 1.01, and 0.81 ° A root-mean-squared deviation (RMSD), respectively, to the segment of the full-length protein that aligns to the isoform. However, this three-dimensional fold is improbable: multiple hydrophobic residues are exposed in a cleft that in the fulllength form of myoglobin, would be occupied by helices A and B. We quantified this effect using the spatial aggregation propensity (SAP) score(10). The bottom row of Fig. 2A depicts the surface of the sequence corresponding to the isoform within the full-length protein, as well as the isoform structure models, colored by the calculated per-residue SAP score. Structure predictions of isoforms from human Prostaglandin E synthase 3 (Fig. 2B), human Caspase-9 (Fig. 2C), and human Nfs1 cysteine desulfurase (Fig. 2D) all share similar trends, where the isoform structure model contains a significant patch of residues with high SAP score. We observed low RMSD to the reference full-length structure, accompanied by high model confidence and increased mean SAP scores across many isoforms (Fig. 2E), indicating both MSA-based and pLM-based models are prone to the error of predicting structures of modified sequences within the context of the full-length protein, countering hypothesis 1.

**Fig. 2.**
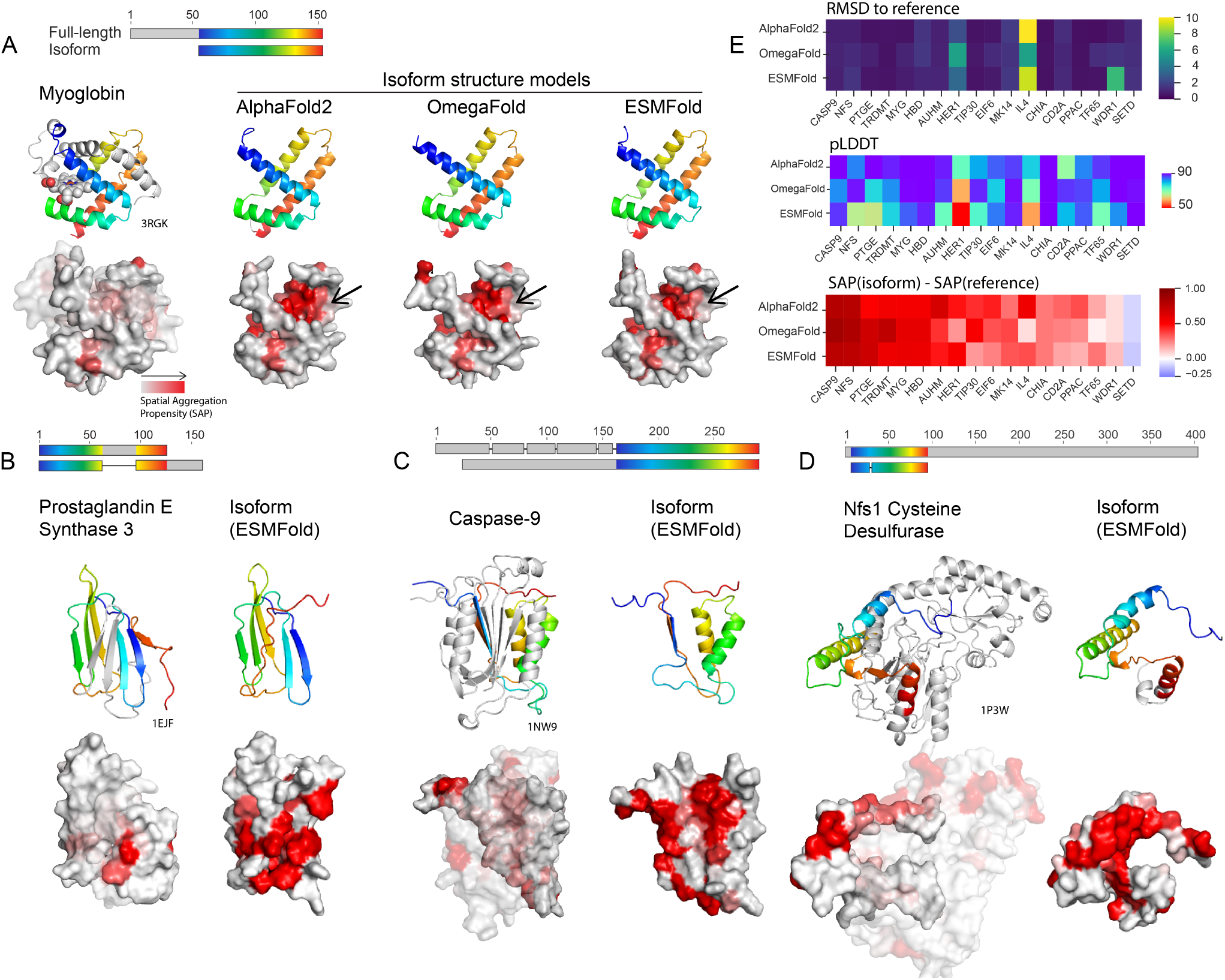
Deep learning structure-based methods predict isoforms as fragments of full-length structures with exposed aggregation-prone residues. A) Top: alignment between human myoglobin (UniProt: P02144) and human isoform Q8WVH6 (UniProt: Q8WVH6). Left: X-ray structure of human myoglobin (PDB: 3RGK) shown with the heme cofactor, and the two missing helices A and B in the isoform colored in grey. Right: predicted structures of isoform Q8WVH6 of Myoglobin from AlphaFold2, OmegaFold, and ESMFold have low root-mean-squared deviation (RMSD) to structure 3RGK (0.49, 1.01, 0.81 A°respectively). Bottom: surfaces of protein fragments corresponding to the isoform sequence. The isoform structure models all have exposed hydrophobic residues corresponding to where helices A and B reside in the full-length structure (indicated with an arrow), quantified here using the spatial aggregation propensity (SAP) score(10). Structure predictions of isoforms from Human Prostaglandin E synthase 3 (B), Human Caspase-9 (C), and Human Nfs1 cysteine desulfurase (D) all share similar trends, where the isoform structure model contains a significant patch of residues with high SAP score. In B-D, the structure model depicted is from ESMFold. (E) For 18 isoforms previously identified in the literature as isoforms where splicing events occur in structured domains, we calculated RMSD to a reference structure of the full-length protein, and the change in average SAP for the isoform fragment in comparison to the sequence aligned in the full-length protein. We found that for AlphaFold2, OmegaFold, and ESMFold, isoform structure models generally had low RMSD to the reference structure, predicted with relatively high pLDDT, along with increased average SAP.

### An unsupervised method of extracting co-evolutionary signal from language models

Following our observations regarding isoforms, we proceeded to further explore how the language model ESM-2, the language model underlying ESMFold (Fig. 3), predicts contacts and how it might be storing coevolutionary information. In ref. (12), the authors first developed a method for contact prediction by supervising training on representations from within the language model, the so-called “Contact Head”. Ref. (4) furthered this work by developing the “Folding Trunk” to predict 3D structure from ESM-2 embeddings. Both of these extensions to the original ESM-2 model were developed using supervised learning on sets of contacts or 3D structures. We wished to develop an approach to evaluate co-evolutionary signal in a completely unsupervised manner, to understand what information the original ESM-2 model, trained only using the unsupervised task of masked language modelling, holds. We formulated the “categorical Jacobian” calculation (Fig. 3) described below toward this end.

**Fig. 3.**
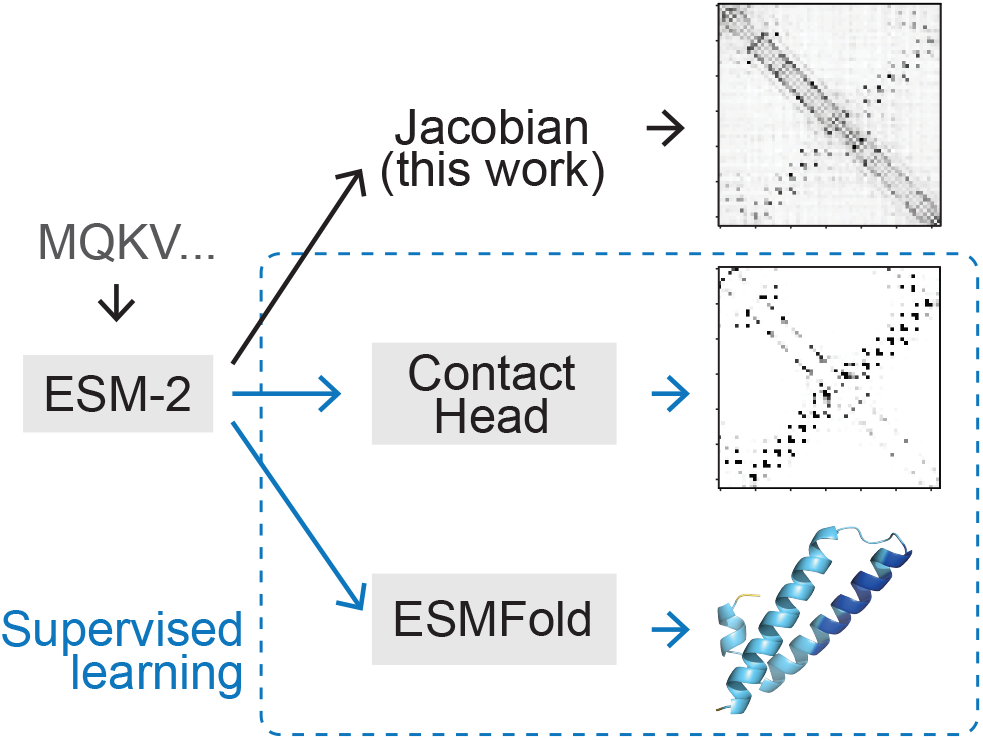
Scheme comparing strategies to extract structure and coevolutionary information from the language model ESM-2. We present an unsupervised “categorical Jacobian” calculation to extract coevolutionary couplings.

For a biological sequence of length *L* with *A* possible tokens (i.e., amino acids for proteins), we extract a set of weights defining the “categorical Jacobian” **J** as follows (illustrated in Fig. 4). We mutate each residue in the sequence to each of *A* possible tokens, and calculate how each of these *L × A* mutations perturbs the probabilities of each amino acid across all positions output by the language model, i.e. the logits, which have shape *L× A*. Accordingly, the shape of the tensor **J** is *L×A×L×A*. Applying the same procedure to a Markov Random Field (MRF)(13–15) or multivariate Gaussian (MG) (16) model results in exactly returning the pairwise coupling tensor **W***L,A,L,A*, and could be also calculated by perturbing the value of the original token, yet we found that in the context of ESM-2, this “categorical” perturbation is critical. In a linear model (MRF or MG), perturbation of any step size returns the same value in the Jacobian (Fig. S1A) (17), yet in ESM-2, a small perturbation to the one-hot encoded input is insufficient to perturb the output (Fig. S1B). We noticed that increasing the step size improves contact map accuracy (Fig. S1C-E) and changing the actual category (amino acid type) results in the best contact accuracy (Fig. S1F). This unsupervised Jacobian method allows us to directly compare pairwise coupling weights from language models to pairwise coupling weights derived from MRF and MG-based models.

**Fig. 4.**
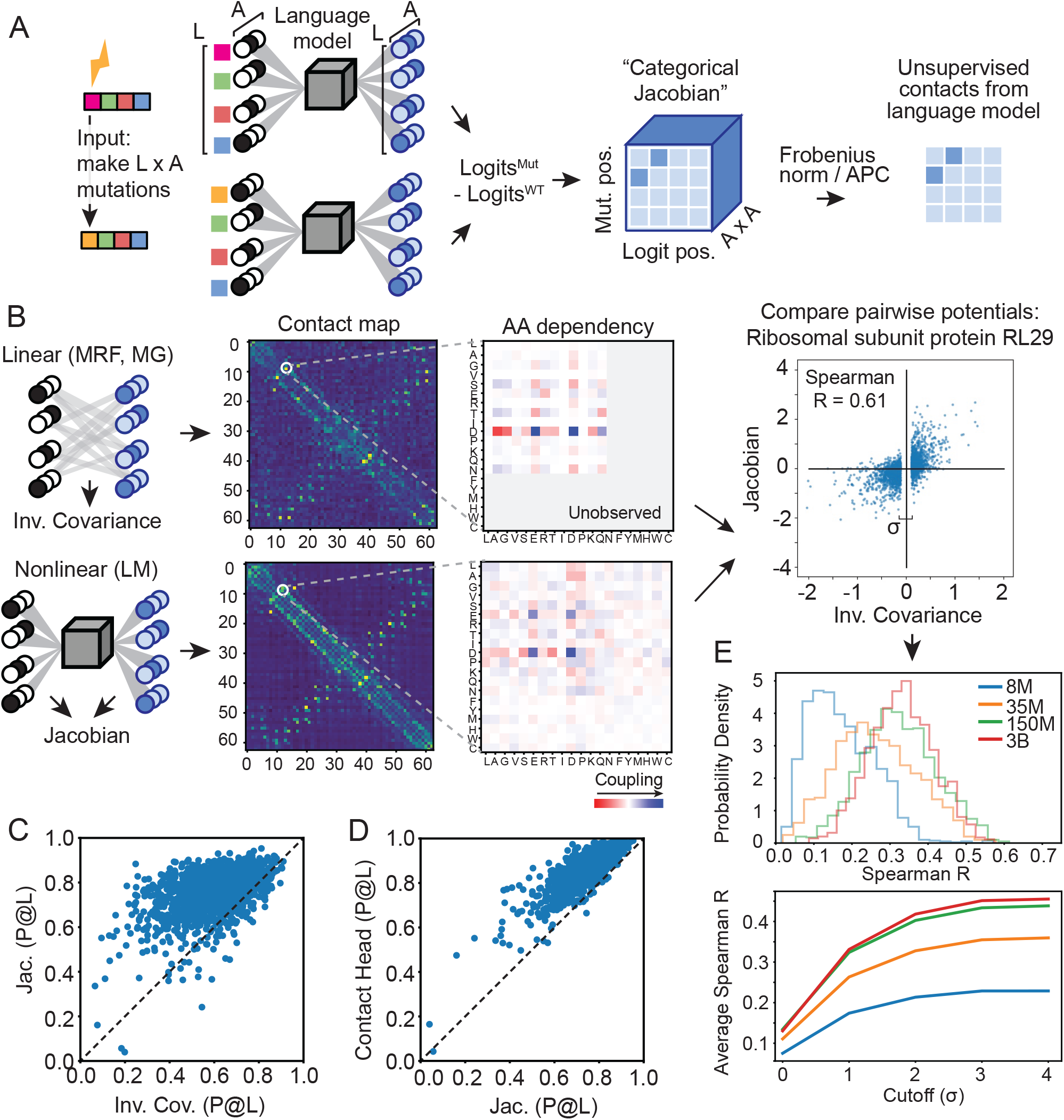
(A) Scheme of the categorical Jacobian calculation. Each residue in a sequence of length L is changed to A different types of amino acids, where A is the size of the alphabet. By computing how the output changes with respect to the input, a matrix of size [L, A, L, A] is obtained. (B) This categorial Jacobian allows for comparing a nonlinear method like ESM-2 and a simple linear method, exemplified here for large ribosomal subunit protein RL29 (UniProt: P0A7M7). We can compare the coevolutionary weights obtained from the a linear model, calculated using inverse covariance, and the categorical Jacobian calculated from ESM-2. (C) Contacts calculated from the categorical Jacobian from ESM-2 outperform the inverse covariance calculation from ref. (11) (Average P@L of 0.61 vs 0.80, respectively). Contacts from the categorical Jacobian are only moderately worse than contacts from the supervised contact prediction head (Average P@L of 0.80 vs 0.87, respectively). (E) Correlation between covariation parameters from linear model and ESM-2 Jacobians increase with model size. Top: Distribution of Spearman correlation coefficients between contacts from linear model and ESM-2 Jacobians. Bottom: average Spearman R, varyingσ cutoff for linear model values close to zero.

With this categorical Jacobian calculation in hand as an unsupervised approach for assessing pairwise coevolutionary weights of pLMs, we next set out to evaluate how these pairwise weights compare to linear models in the task of contact prediction, as well as the supervised “Contact Head” that was trained on top of ESM-2 embeddings. From our Jacobian tensor, we can calculate a predicted contact map of size *L × L* analogously to MRFs and MGs (see Methods). Fig. 4B depicts an example comparison of pairwise coevolutionary weights for the large ribosomal subunit protein RL29 calculated with 2 methods: on the left, using a multivariate Gaussian approach inferred from an MSA for the family, and on the right, from the categorical Jacobian of ESM-2 with 3 billion parameters. The top row depicts summed contact weights from both methods and the bottom row depicts an example 20x20 set of weights corresponding to pairwise amino acid dependencies for a given pairwise contact, demonstrating striking visual similarities between the two methods. Analogous sets of weights for other ESM-2 model sizes are depicted in Fig. S2.

We compared the accuracy of contacts predicted with a standard linear model for pairwise couplings (11, 18) or predicted with the categorical Jacobian of the ESM-2 3-billion-parameter model, quantifying accuracy via precision of the *L* top-weighted contacts (precision@L). We used the 3-billion-parameter model because it showed similar performance to the 15-billion-parameter model (4). The categorical Jacobian calculation demonstrated improved performance at predicting contacts than the linear model across our dataset of 1431 proteins (Fig. 4C, average accuracy of 0.80 and 0.61, respectively). Furthermore, contacts from the categorical Jacobian were not that much worse than contact accuracy from the supervised contact head (Fig. 4D), average accuracy of 0.80 and 0.87, respectively).

Next, we were curious how similar the actual underlying weight matrices were between these two methods, i.e., a linear model and the ESM-2 categorical Jacobian calculation. Estimating a linear model involves fitting *L × A × L × A* parameters for each family, which is very likely overdetermined, and many of the weights are driven to zero. We assessed the correlation at different cutoffs of removing weights closest to zero (see Methods). In our benchmark of 1431 proteins, we found that the correlation between pairwise coupling weights from ESM-2 and from a linear model increased with the size of the ESM-2 model (Fig. 4E), with performance plateauing at the 150-million to 3-billion parameter model sizes.

### Language models predict structures by looking up segment pairings

Given that we could calculate a Jacobian of ESM-2 that contained coevolutionary signal rivalling the information predicted by the supervised Contact Head, we wished to more thoroughly investigate the mechanism of precisely how the Contact Head predicts contacts given an input sequence. We tested what information is most used in contact prediction by monitoring contact prediction when information from various sequence locations is masked. We compared unmasking flanking residues next to a contact, randomly unmasking any residues, and randomly unmasking any residues more than 30 residues away from the contact (Fig. 5A). We hypothesized that a model that had memorized complete folds would be able to recover contacts by randomly unmasking residues and that a model that had memorized local motifs would better be able to recover contacts by gradually unmasking residues next to the contact in question. Testing these unmasking strategies revealed that ESM-2 more rapidly recovered contacts by unmasking flanking regions, with 50% being recovered with approximately 30 residues unmasked on each side, and contact recovery from flanking unmasking being roughly twice as effective as randomly unmasking (Fig. 5B). We observed a striking step-function type behavior in how ESM-2 uncovered contacts while unmasking flanks, an example of which is shown for the ATP-dependent dethiobiotin synthetase BioD 1 protein (1BYI) (19). ESM-2 shifts from not predicting the contact using 10 flanking residues on each side, to complete contact recovery at 20 residues (Fig. 5C-D).

**Fig. 5.**
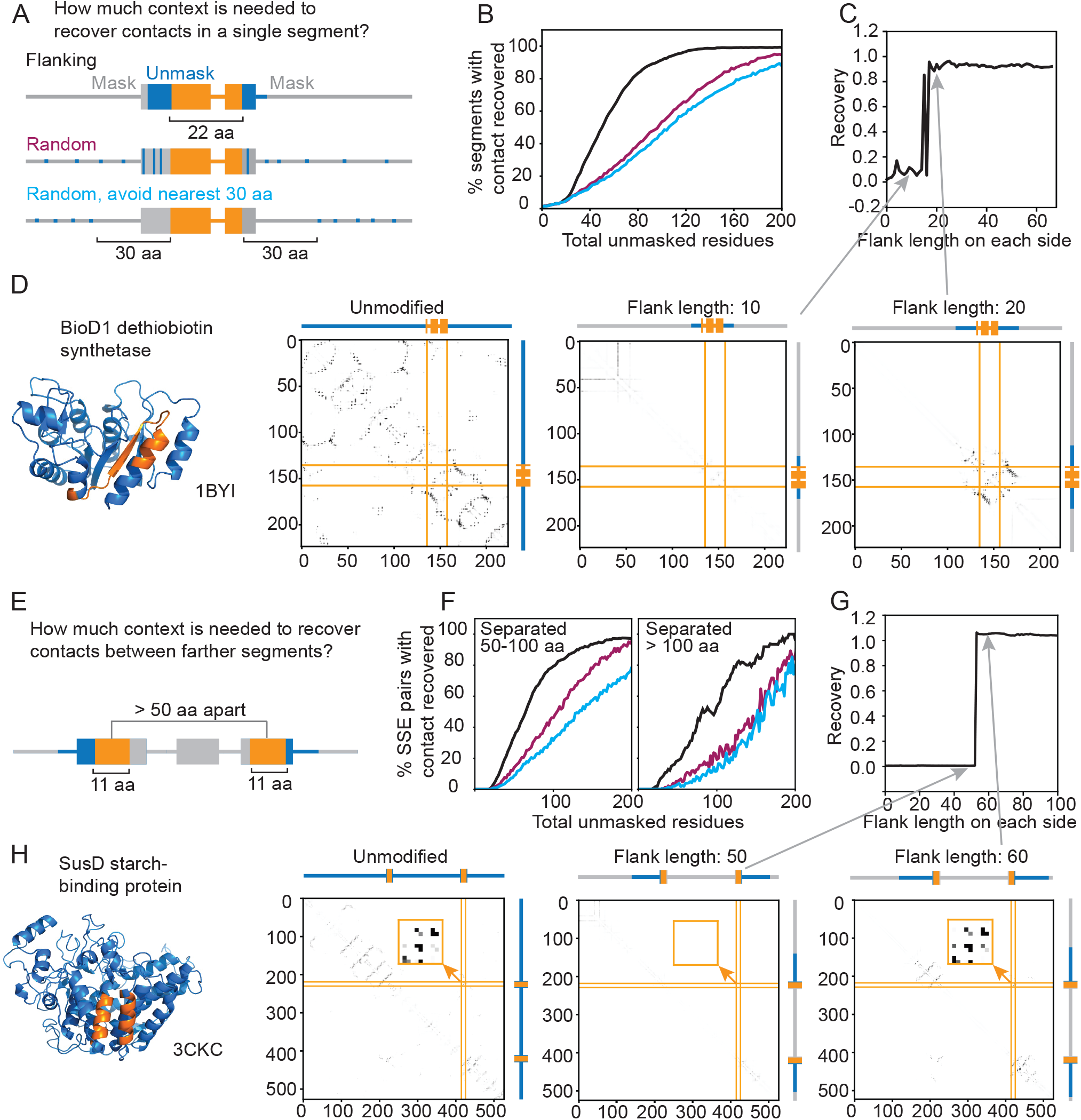
(A) Scheme depicting the single segment contact recovery experiment. (B) The percentage of segments with contact recovered at different numbers of unmasked residues showed that the contact within most single segments was recovered after extending the segment each way for 30 aa, and random unmasking required 100-120 unmasked residues for contact recovery. (C-D) In ATP-dependent dethiobiotin synthetase BioD 1 protein (PDB 1BYI), the contact within a single segment was recovered by extending each side outward for 20 residues. (E) Scheme depicting the SSE pair contact recovery experiment. (F) The percentage of SSE pairs with contact recovered at different flanking lengths showed that the contact of most SSE pairs was recovered after extending each SSE outwards for 30-40 aa (60-80 aa unmasked in total), while random unmasking required 100-160 unmasked residues. (G-H) In starch-binding protein SusD (PDB 3CKC) the contact between the SSE pair with centers at residue 225 and 421 was recovered by extending both SSEs outward for 60 residues.

We found similar trends when analyzing how much context ESM-2 needs to recover contacts between more distant secondary structure elements (SSEs). We took two 11residue segments from a pair of interacting SSEs, with centers separated by at least 50 residues, and masked the rest of the protein. Then, we gradually unmasked more flanking regions on the outer sides of the segments (Fig. 5E) and monitored the contact recovery. We found that 50% of the SSE pairs’ contacts were restored with a flanking length of 30 or 40 residues (Fig. 5F) for pairs separated by 50-100 or more than 100 residues, respectively. The contact recovery from flanking unmasking is again roughly twice as effective as random unmasking (Fig. 5F). We also observe the same stepwise behavior in contact recovery, exemplified in starch-binding protein SusD (PDB 3CKC) (20) in (Fig. 5G-H). The contact between two α-helices centered at residue 225 and residue 421 appears in the LM contact map when the flanking region reaches approximately 60 residues (Fig. 5GH). Another example of this is given in Fig. S3 for a contact between two β-strands in the ATPase region of topoisomerase II (PDB 1PVG) (21).

We further explored the effects of tokens and positional embedding on contact recovery. During ESM-2 training, BOS and EOS tokens are used to indicate the start and end of the protein for the model to distinguish a full-sized protein from a cropped one. Replacing these tokens at the start and end of each protein with a mask token reduced the amount of context required for contact recovery (Fig. S4). Instead of masking, positions can also be removed and the residue index can be offset to account for the removal (Fig. S5A). We find removal+offset to result in only 10% decrease in performance, suggesting the model is able to perform segment pairing without space needed for in-painting or inferring the full fold (see Methods, Fig. S5B). Notably, we also found that sheet-sheet interactions demanded less context compared to helix-helix interactions (Fig. S6). This is consistent with the notion that β-strands are frequently composed of fewer residues than α-helices, and that β-strands tend to form stable β-sheets. This difference in context needed for recovery for helices and strands was much more pronounced when using offset compared to masking (Fig. S6).

## Discussion

The development of protein language models (pLMs) has brought significant excitement into the field of protein structure prediction. Some have wondered if pLMs have finally solved the “protein folding problem”, given their accurate structure prediction from single sequences and no supplied co-evolutionary signal in an input multiple sequence alignment(2). This should have been quickly debunked, as the accuracy of models was found to be highly correlated to the number of related proteins in the training set(3, 4), indicating that the models store evolutionary information in their parameters, but precisely how has been unclear.

A clue for how ESM-2 might be storing coevolutionary information came via a consistent error we encountered in the predicted structures of isoforms, which we found were consistently predicted to fold to fragments matching their structure context within the full-length proteins, but which left non-physical patches of hydrophobic residues exposed. We figured if the model learned protein folding and not simply looked up evolutionary statistics, it should be able to model a more-likely unfolded conformation. Our results caution against assuming pLMs as oracles of protein properties without consideration of potential adversarial and out-of-distribution behaviors. Notably, AF2 is prone to this error as well. As of January 2024, we identified one such erroneous structure, Myoglobin isoform CHS.35702.2 predicted with pLDDT 94.3 (Fig. S7), from the isoforms we analyzed in the “CHESS human protein structure database” public database of AF2 isoform predictions (22). A clear limitation of this study is that we do not have experimental evidence for the actual in vitro structure landscapes of these isoform examples.

Motivated to develop a framework to assess coevolutionary signal within language models, we developed a general calculation to calculate a “categorical Jacobian” of a pLM for a given sequence. The values of the categorical Jacobian can be directly compared to the pairwise weights of a Markov Random Field(13–15) or multivariate Gaussian model(16) calculated for a given MSA, approaches which have long been used to assess coevolutionary couplings in protein families. We found that the weights derived via this categorical Jacobian achieved accuracy in predicting contacts not that much lower than those calculated with the Contact Head, trained with supervised learning (mean accuracy 0.80 vs. 0.87, respectively).

We were curious if we could detect patterns in how ESM-2 uses coevolutionary information to predict contacts. We tested unmasking residues in various patterns surrounding contacts, which revealed that the model best recovers contacts by gradually unmasking residues next to the contact in question comparing to random unmasking. This suggests that pLMs learned statistics of motif pairings. We suggest that this relationship can be roughly represented as:

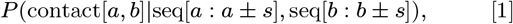

where *s* is approximately 20-40 residues. Without this context, the pLMs are unable to correctly predict the interaction between fragments.

Our analysis does not completely rule out that pLMs have learned the concept of full folds, since the continuous segment unmasked in the flanking region might have helped the model to match to full proteins. Nevertheless, our results underscores that the information of the full fold is not required for the model to function.

Storing the co-evolutionary statistics^*^ of all known protein families in UniProt (roughly 20,000), assuming an average length of 256, would require

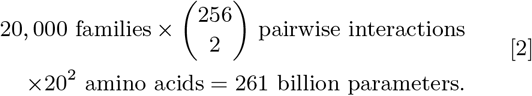

If we assume each position makes at most 4 contacts – 2 sequence neighbors and roughly 2 long-range contacts – this corresponds to

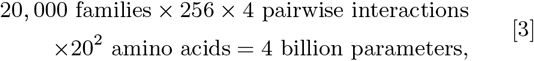

which is the same order of magnitude at which ESM-2 models start to taper off in their improvement (roughly 3 billion parameters)(12). A model that segments a protein into common motifs, as our work suggests pLMs are doing, offers a clear route to compression. A downside of such compression is that within-family evolutionary effects such as multiple stable conformations are inaccurately predicted by ESM-2 (see SI), a clear area for future improvement.

In summary, our work has demonstrated how a fundamentally powerful unsupervised learning approach – that of masked language modelling – enables storing coevolutionary statistics agnostically for thousands of protein families. Although they have not yet reached the ability to directly model the physics of protein folding, we anticipate that this research and other ongoing interpretability studies will shed light in how we might actually use deep learning to approximate the fundamentals of biophysics.

## Data and code availability

The dataset of 18 isoforms and scripts to perform analysis are available at https://github.com/HWaymentSteele/Isoforms benchmark 2024. The code for categorical Jacobian and contact prediction analyses is available at https://github.com/zzhangzzhang/pLMs-interpretability. The modified positional embedding version of ESM-2 and ESMFold are available at https://github.com/garykbrixi/esm_gap_distance.

## Materials and Methods

### Isoform dataset curation and analysis

We collected examples of isoforms identified in previous literature(6–9) as cases where splicing would disrupt ordered domains, along with associated structures. For each, we identified a corresponding isoform and full-length protein in UniProt. We predicted structure models in AlphaFold2(1) using ColabFold(23); OmegaFold(2) using the OmegaFold notebook available at https://colab.research.google.com/github/sokrypton/ColabFold/blob/main/beta/omegafold.ipynb; and ESMFold(4) using the ESMFold server available at https://esmatlas.com/resources?action=fold.

We calculated the Surface Aggregation Propensity (SAP) score of isoform models using the “per res sap.xml” script from the supporting materials of ref. (24). We calculated root-mean-squared deviation (RMSD) between structure models of isoforms and the full-length experimental structure in PyMOL(25) for α-carbons for aligned regions. Aligned regions were manually determined from alignments of each isoform to each full-length sequence using the “global alignment with free end gaps” setting and the BLOSUM62 matrix in the Geneious Prime software.

### Dataset for model comparison and contact recovery

We obtained 2245 structures from the GREMLIN Coevolution predictions database for PDB EXP with more than 1000 sequences in MSA (14). Similar structures were filtered based on a TMalign (26) score exceeding 0.5. We used proteins with a length from 200 to 600 amino acids to ensure a similar size across proteins and enough length for exploring the effect of flanking region (Fig. S8A). We selected only proteins that were missing fewer than 50 residues in the structure to ensure the majority of residues are present in the experimental structure. With these filtering steps, we end up with a dataset of 1431 proteins in total.

### Weights and contact maps from a linear model

To calculate pairwise coupling weights and contact maps from MSAs via a linear model, we use the inverse covariance method presented in (11). Dauparas et al. demonstrated that Markov Random Field (MRF) and multivariate Gaussian (MG) models can be mapped to the same graphical model representation. The main difference is that MRFs consider biological sequences as categorical, and MGs approximate them as continuous variables. Estimating a set of pairwise coupling weights **W***L,A,L,A* for either then depends on the loss function used. Dauparas et al. demonstrated empirically that the following estimation, which derives from a mean-squared-error loss for an MG formalism, performs comparably to cross-entropy loss in an MRF, as used in GREMLIN(13) and other models, but with substantially less compute. We defer the reader to (11) for the complete derivation.

In brief, a set of sequences in an MSA is one-hot-encoded and written in the form *X ∈* ℝ^*N×LA*^, where *N* is the number of sequences, *L* is the sequence length, and *A* is the number of letters available in the alphabet (*A* = 20 for proteins). The pairwise coupling weights **W** are calculated as

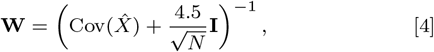

where 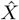 is the mean-centered MSA, i.e.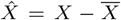, and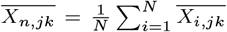 for all *n*. The 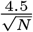 term is introduced for shrinkage and is empirically estimated in (11). Note that the expression for **W** above is technically the weight matrix minus the identity matrix. This is expressed in (11) as 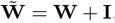, but this makes no difference in calculation.

We calculate a contact matrix *C ∈* ℝ^*LxL*^ with entries *c*_*ij*_ from the above tensor **W** using

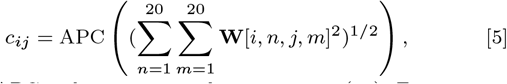

where APC is the average product correction (27). For a matrix in ℝ^*L×L*^ comprised of entries *m*_*ij*_, we calculate the APC as

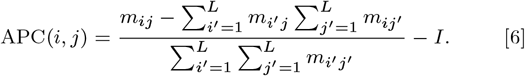

### Language Model Contact Map via Jacobian

To calculate the categorical Jacobian of a language model, each of *L* positions in a sequence is mutated to all *A* possible tokens (for proteins, *A* = 20) and input into the language model to predict the resulting logits across the entire sequence, where the logits are shaped *L × A*. The difference between the logits of the original sequence and the logits of the mutated sequences was calculated to get the Jacobian matrix.

Written formally, we define the categorical Jacobian **J** for a protein language model *f* (*X*), which accepts as its input a protein sequence *X* with length *L* and alphabet size *A*, as

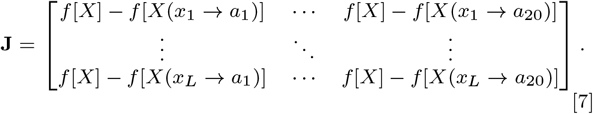

Above, *f* [*X*] is the original logits output by the language model, a matrix of logits with size *L ×A. f* [*X*(*x*_*i*_ *→ a*_*n*_)] represents the logits returned when position *i* has been mutated to token *an*. The Jacobian is therefore a tensor with size [*L, A, L, A*]. **J** is mean-centered and symmetrized. We obtain a contact map from **J** analogously to equations 5 and 6. It can be shown that applying the same categorical Jacobian operation to an MRF or MG model will return the pairwise weights matrix **W**.

To evaluate the correlation between the pairwise couplings from ESM-2 models and the couplings from a linear model, we first selected the top-weighted *L* inter-residue contacts from the contact map calculated from the linear model (following average product correction). This results in Lx20x20 weights from both the linear model couplings (**W**) and the ESM-2 jacobian **J**. We expect many of these weights to be close to zero and not meaningful, so we calculate Spearman correlation over a range of cutoffs filtering values close to zero.

If 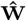 is this reduced mean-centered set of couplings from the linear model, and 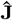 is the reduced mean-centered ESM-2 jacobian, we calculate the Spearman correlation over the subset of 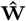and 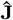 whose absolute value is greater than *b* standard deviations of 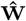, where *b ∈* [0, 4]. The Spearman correlation is calculated in SciPy (28).

### Recovery of contact with increasing flanking region

For the extraction of secondary structures, we first predicted structures from sequences with ESMFold and then used PyDSSP (29). We chose pairs of secondary structure elements (SSEs) with centers separated by at least 50 amino acids, because the contact probability derived from the GREMLIN dataset showed correlations diminished beyond 40 residues (Fig. S8B). The contact recovery between the SSE pairs was calculated via

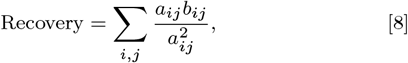

where *a*_*ij*_ and *b*_*ij*_ are the contact probabilities corresponding to positions *i, j* from the LM contact of the original and masked sequences, respectively. When the score was higher than 0.5, we regarded it as a recovery of contact.

### Modification of the rotary position embeddings

We modified the rotary position embeddings (RoPE) (30) to allow modifying the positions of the amino acids without adding additional tokens between them. We achieved this by extending the length of the sine and cosine tables and retrieving the desired index for each amino acid from the sine and cosine tables. This way, the total length of the transformer input and operations stayed the same, but the RoPE index of each query and key corresponded to the offsets.

## ACKNOWLEDGMENTS

We thank Ramya Rangan, Sirui Liu,and members of the Ovchinnikov lab for useful discussion. H.K.W.S. acknowledges funding from the Jane Coffin Childs foundation and HHMI. Z.Z. and M.D.P. acknowledges the Swiss National Science Foundation. D.K. acknowledges funding from HHMI. S.O. and H.W. were supported by NIH [DP5OD026389] and NSF [MCB2032259].

## Footnotes

* Storing sequence conservation requires (20,000 protein families *×* 256 sequence positions *×* 20 amino acids) = 100 million parameters for any model.

## References

1. J Jumper, et al., Highly accurate protein structure prediction with AlphaFold. Nature 596, 583–589 (2021).

2. R Wu, et al., High-resolution de novo structure prediction from primary sequence, (Bioinformatics), preprint (2022).

3. R Chowdhury, et al., Single-sequence protein structure prediction using language models from deep learning, (Bioinformatics), preprint (2021).

4. Z Lin, et al., Evolutionary-scale prediction of atomic-level protein structure with a language model. Science 379, 1123–1130 (2023).

5. M Baek, et al., Accurate prediction of protein structures and interactions using a three-track neural network. Science 373, 871–876 (2021).

6. F Birzele, G Csaba, R Zimmer, Alternative splicing and protein structure evolution. Nucleic acids research 36, 550–558 (2008).

7. ML Tress, B Bodenmiller, R Aebersold, A Valencia, Proteomics studies confirm the presence of alternative protein isoforms on a large scale. Genome Biol. 9, R162 (2008).

8. S Light, A Elofsson, The impact of splicing on protein domain architecture. Curr. opinion structural biology 23, 451–458 (2013).

9. F Pozo, et al., Assessing the functional relevance of splice isoforms. NAR Genomics Bioinforma. 3, lqab044 (2021).

10. N Chennamsetty, V Voynov, V Kayser, B Helk, BL Trout, Design of therapeutic proteins with enhanced stability. Proc. Natl. Acad. Sci. 106, 11937–11942 (2009).

11. J Dauparas, et al., Unified framework for modeling multivariate distributions in biological sequences (2019) arXiv:1906.02598 [q-bio].

12. A Rives, et al., Biological structure and function emerge from scaling unsupervised learning to 250 million protein sequences. Proc. Natl. Acad. Sci. 118, e2016239118 (2021).

13. S Balakrishnan, H Kamisetty, JG Carbonell, SI Lee, CJ Langmead, Learning generative models for protein fold families. Proteins: Struct. Funct. Bioinforma. 79, 1061–1078 (2011).

14. H Kamisetty, S Ovchinnikov, D Baker, Assessing the utility of coevolution-based residue–residue contact predictions in a sequence- and structure-rich era. Proc. Natl. Acad. Sci. 110, 15674–15679 (2013).

15. M Ekeberg, C Lövkvist, Y Lan, M Weigt, E Aurell, Improved contact prediction in proteins: using pseudolikelihoods to infer potts models. Phys. Rev. E 87, 012707 (2013).

16. C Baldassi, et al., Fast and accurate multivariate gaussian modeling of protein families: predicting residue contacts and protein-interaction partners. PloS one 9, e92721 (2014).

17. D Marshall, et al., The structure-fitness landscape of pairwise relations in generative sequence models. BioRxiv pp. 2020–11 (2020).

18. J Yang, et al., Improved protein structure prediction using predicted interresidue orientations. Proc. Natl. Acad. Sci. 117, 1496–1503 (2020).

19. T Sandalova, G Schneider, H Käck, Y Lindqvist, Structure of dethiobiotin synthetase at 0.97 A°resolution. Acta Crystallogr. Sect. D Biol. Crystallogr. 55, 610–624 (1999).

20. NM Koropatkin, EC Martens, JI Gordon, TJ Smith, Starch Catabolism by a Prominent Human Gut Symbiont Is Directed by the Recognition of Amylose Helices. Structure 16, 1105–1115 (2008).

21. S Classen, S Olland, JM Berger, Structure of the topoisomerase II ATPase region and its mechanism of inhibition by the chemotherapeutic agent ICRF-187. Proc. Natl. Acad. Sci. 100, 10629–10634 (2003).

22. MJ Sommer, et al., Structure-guided isoform identification for the human transcriptome. eLife 11, e82556 (2022).

23. M Mirdita, et al., Colabfold: making protein folding accessible to all. Nat. methods 19, 679–682 (2022).

24. L Cao, et al., Design of protein-binding proteins from the target structure alone. Nature 605, 551–560 (2022).

25. WL DeLano,, et al., Pymol: An open-source molecular graphics tool. CCP4 Newsl. Protein Crystallogr 40, 82–92 (2002).

26. Y Zhang, TM-align: a protein structure alignment algorithm based on the TM-score. Nucleic Acids Res. 33, 2302–2309 (2005).

27. SD Dunn, LM Wahl, GB Gloor, Mutual information without the influence of phylogeny or entropy dramatically improves residue contact prediction. Bioinformatics 24, 333–340 (2008).

28. P Virtanen, et al., Scipy 1.0: fundamental algorithms for scientific computing in python. Nat. methods 17, 261–272 (2020).

29. W Kabsch, C Sander, Dictionary of protein secondary structure: Pattern recognition of hydrogen-bonded and geometrical features. Biopolymers 22, 2577–2637 (1983).

30. J Su, et al., RoFormer: Enhanced Transformer with Rotary Position Embedding. (2021) Publisher: arXiv Version Number: 4.

