## Supporting Information for "Protein language models learn evolutionary statistics of interacting sequence motifs"

**This PDF file includes:**

Supporting text
Figs. S1 to S10
SI References

### Supporting Information Text

**ESM2 predicted same fold for two KaiB sequences with different experimental structures.** KaiB is a metamorphic protein that adopts two distinct states as part of its function. We extracted contacts for constructs demonstrated in vitro to be stabilized for both states with ESM2. While experimental results showed that each sequence favors one state over another, ESM2 predicted that both sequences fold to the thermodynamically unfavorable fold-switched (FS) state (Fig. S9). ESM2 showed the same failure as the previous MSA based methods such as MSA transformer and AF2 with default settings (1).

**ESM2 predicted same interchain contacts for bacterial response regulator subfamilies with diverse interchain contacts.** Three bacterial response regulator subfamilies were shown to have diverse interchain contacts between their homomeric interfaces despite having similar intrachain contacts (2). This provides an incisive example of a potential pitfall of evolutionary models, where the properties of the protein family lead to a prediction which does not match the property of the sequence.

We observed that the ESM2 predictions for all three sequences capture similar fractions of the true interchain contacts (Fig. S10). Furthermore, the predicted contacts of the three sequences were not closer to the true interchain contacts of that sequence compared to the interchain contacts of the other sequences.

This type of failure is similar to the outputs of MSA-based methods, which did not differentiate different subfamilies if the aligned sequences include sequences from many different subfamilies with different properties.

**Interchain contact comparison for bacterial response regulator subfamilies.** We calculated the interchain contacts from the experimental structures (contact defined as $< 10$ Å) and extracted the ESM2 contact maps for each of the sequences. The outputs of the ESM2 contacts and PDB structure-based contacts were all aligned using US-align, removing all gap positions. The intrachain contacts of all structures were removed, leaving only homomeric interactions unique to that structure.

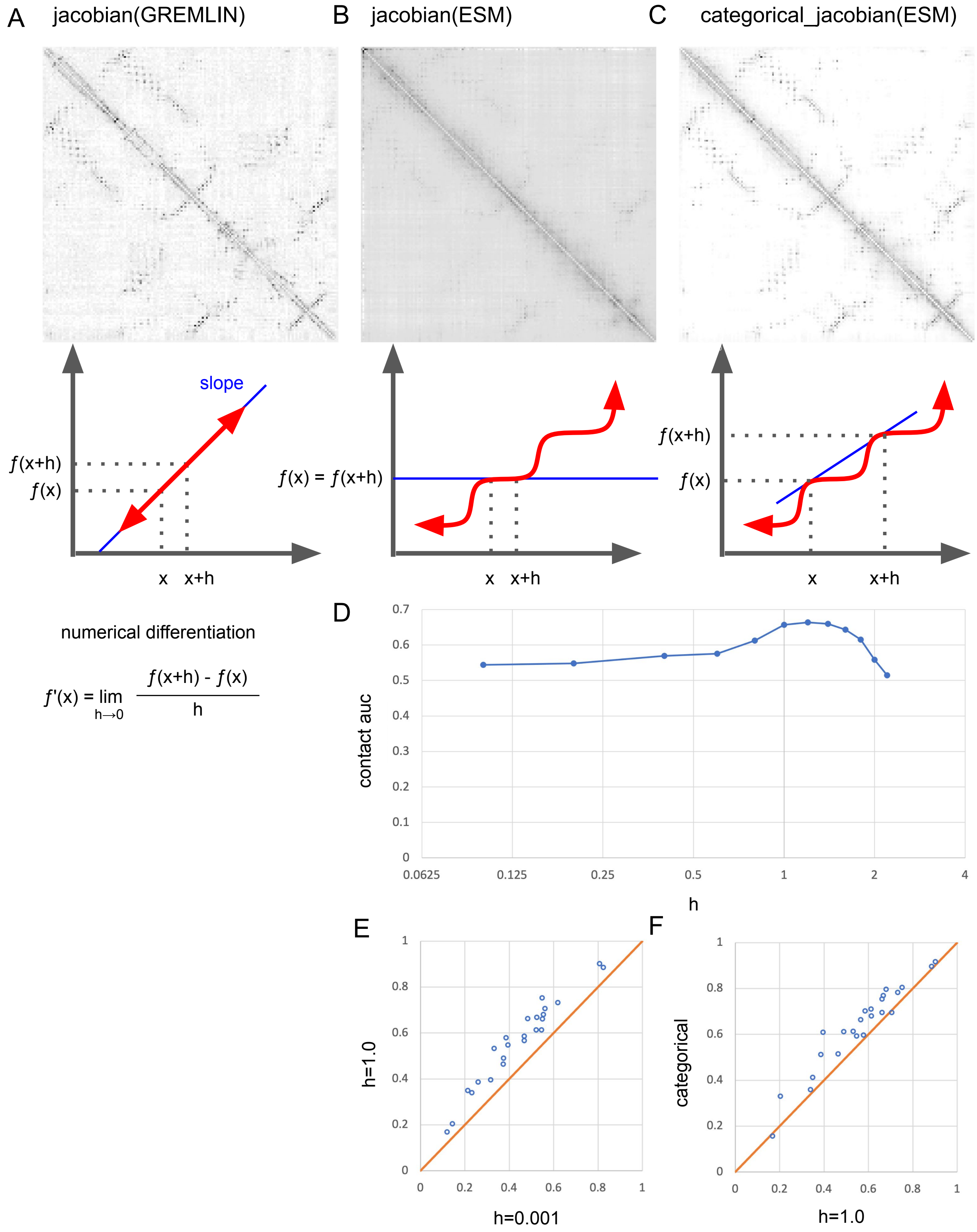


**Fig. S1.** Categorical Jacobian allows capturing non-linear relationships between the input and output of protein Language models. (A) for linear model such as GREMLIN, perturbation of any step size returns the same slope. (B) for non-linear model, a small perturbation might not be enough to change the output. (C,D,E) increasing the step size improves contact map accuracy. (F) flipping the actual category results in best contact accuracy.

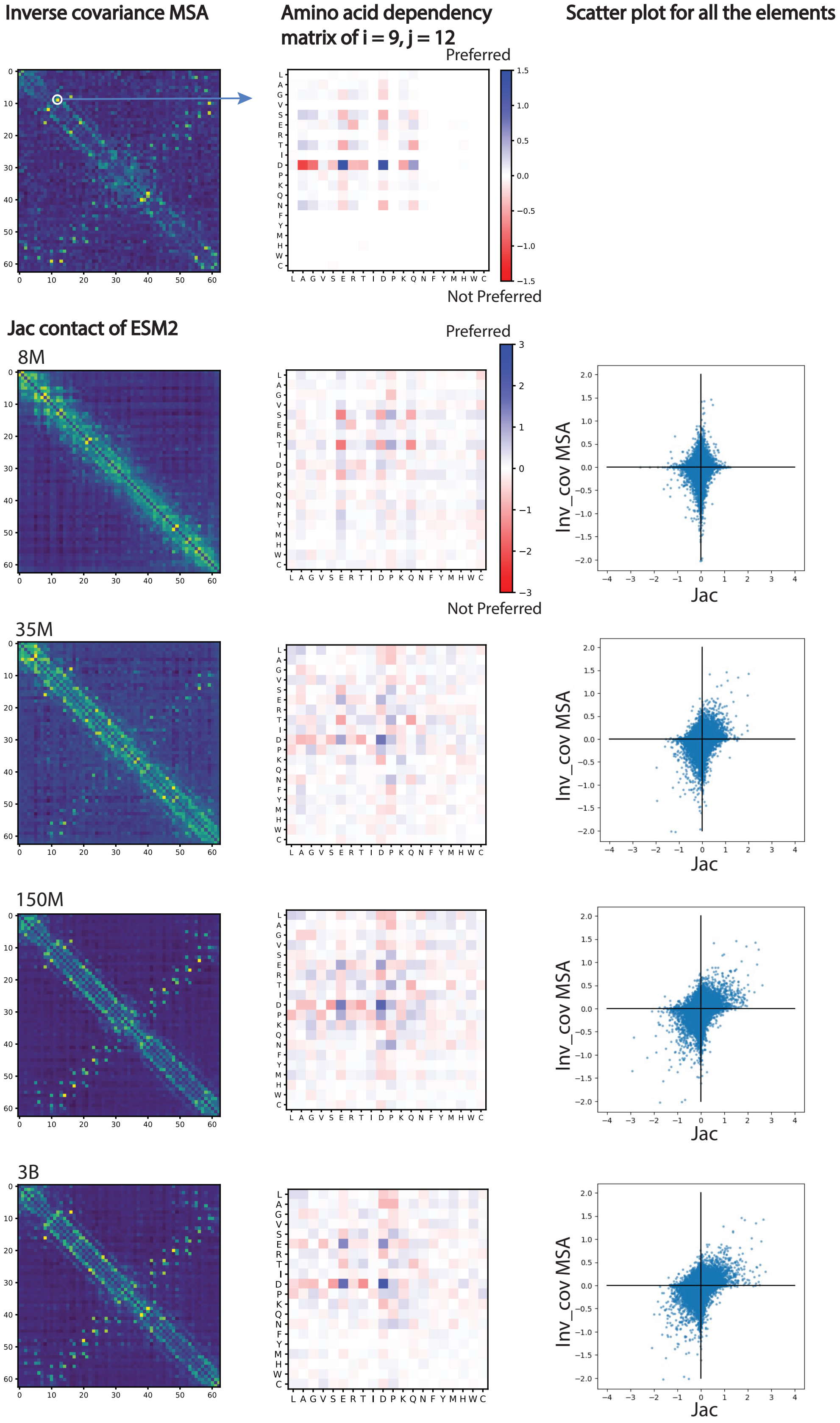


**Fig. S2.** The Jacobian contact maps, amino acid dependency plot at i = 9 and j = 12, and the scatter plot for all elements of ESM models of different sizes showed that larger ESM2 models captured evolution information better

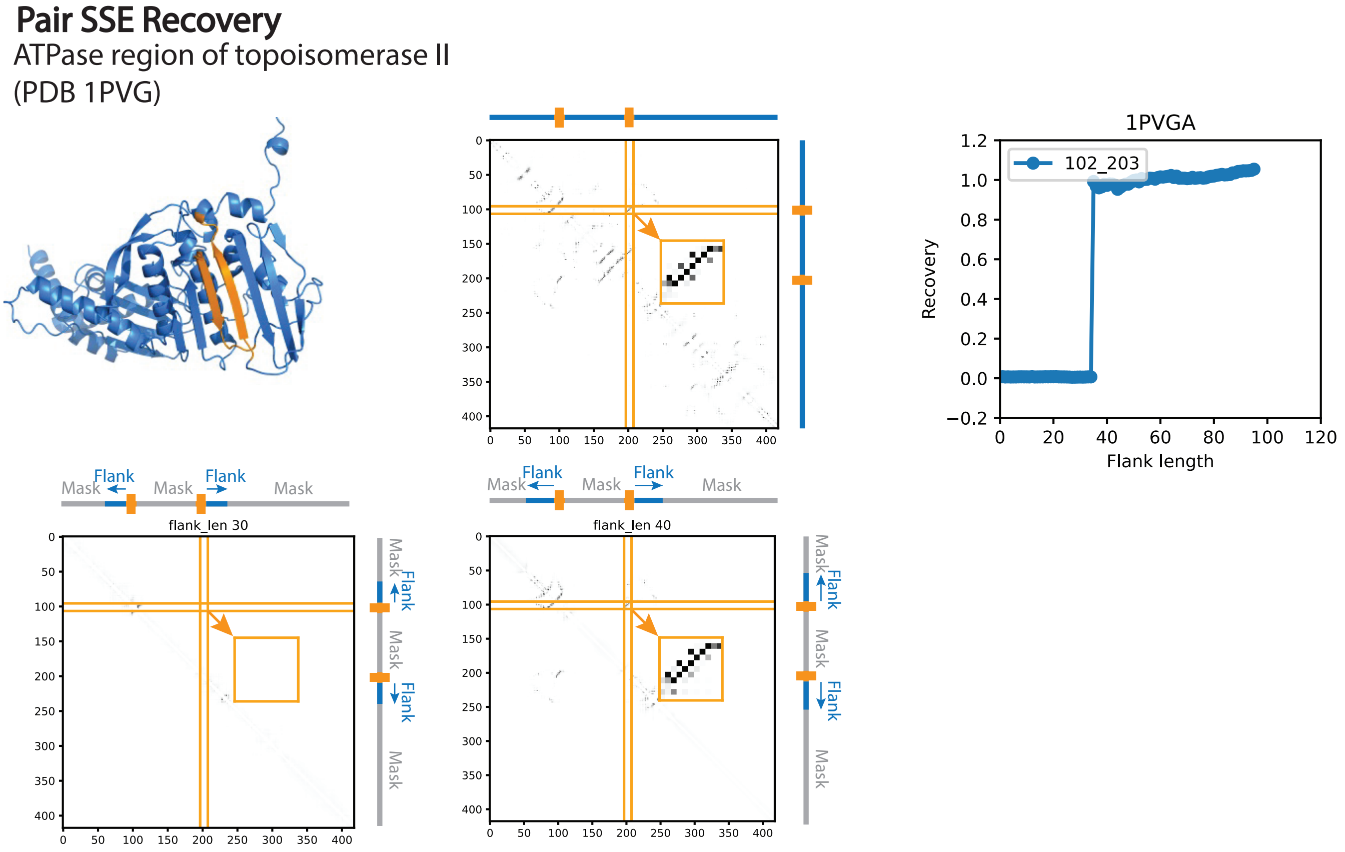


**Fig. S3.** Contact recovery of the ATPase region of topoisomerase II (PDB 1PVG) for a segment pair with centers at residue 102 and 203. The recovery was achieved with around 40 residues.

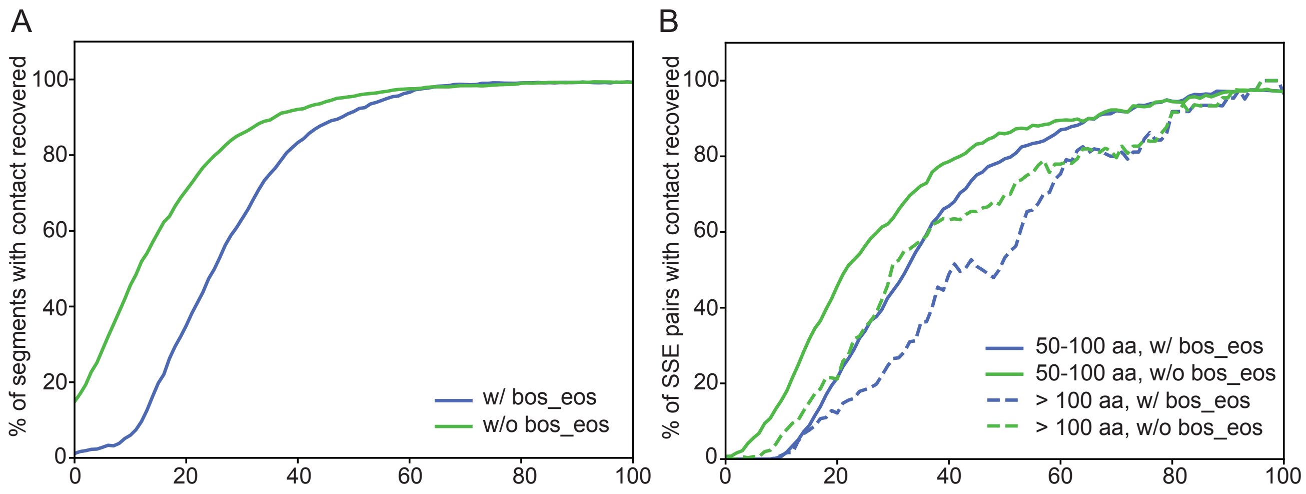


**Fig. S4.** The recovery of contacts required less flanking region without BOS and EOS compare to with BOS and EOS for both (A) single segment and (B) segment pairs.

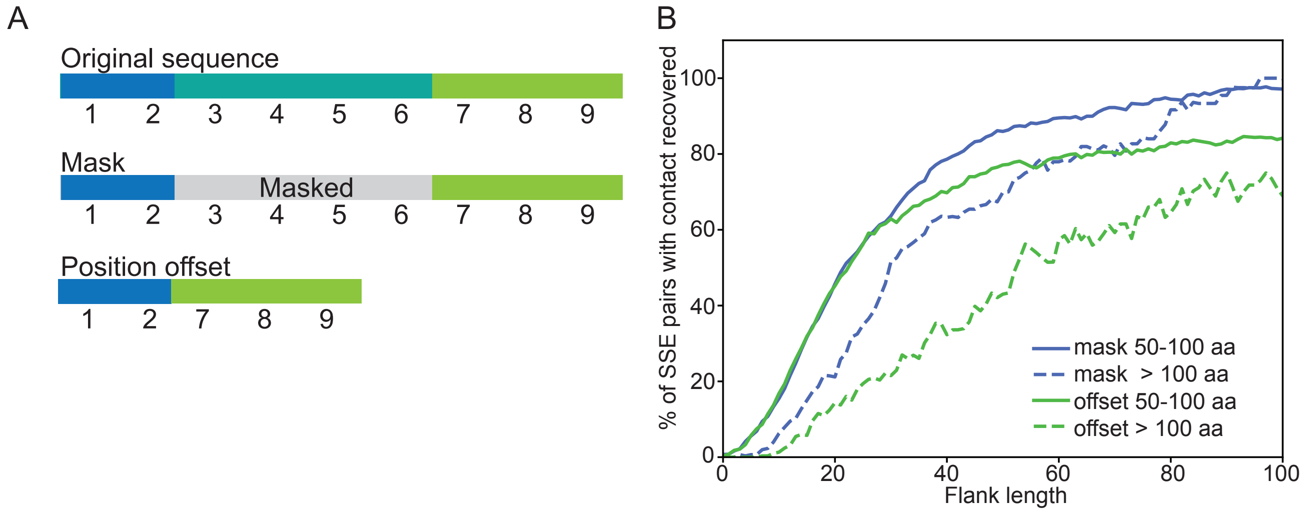


**Fig. S5.** (A) Scheme depicting masking and position offset. (B) Masking resulted in higher recovery of SSE pair contacts compared to offset.

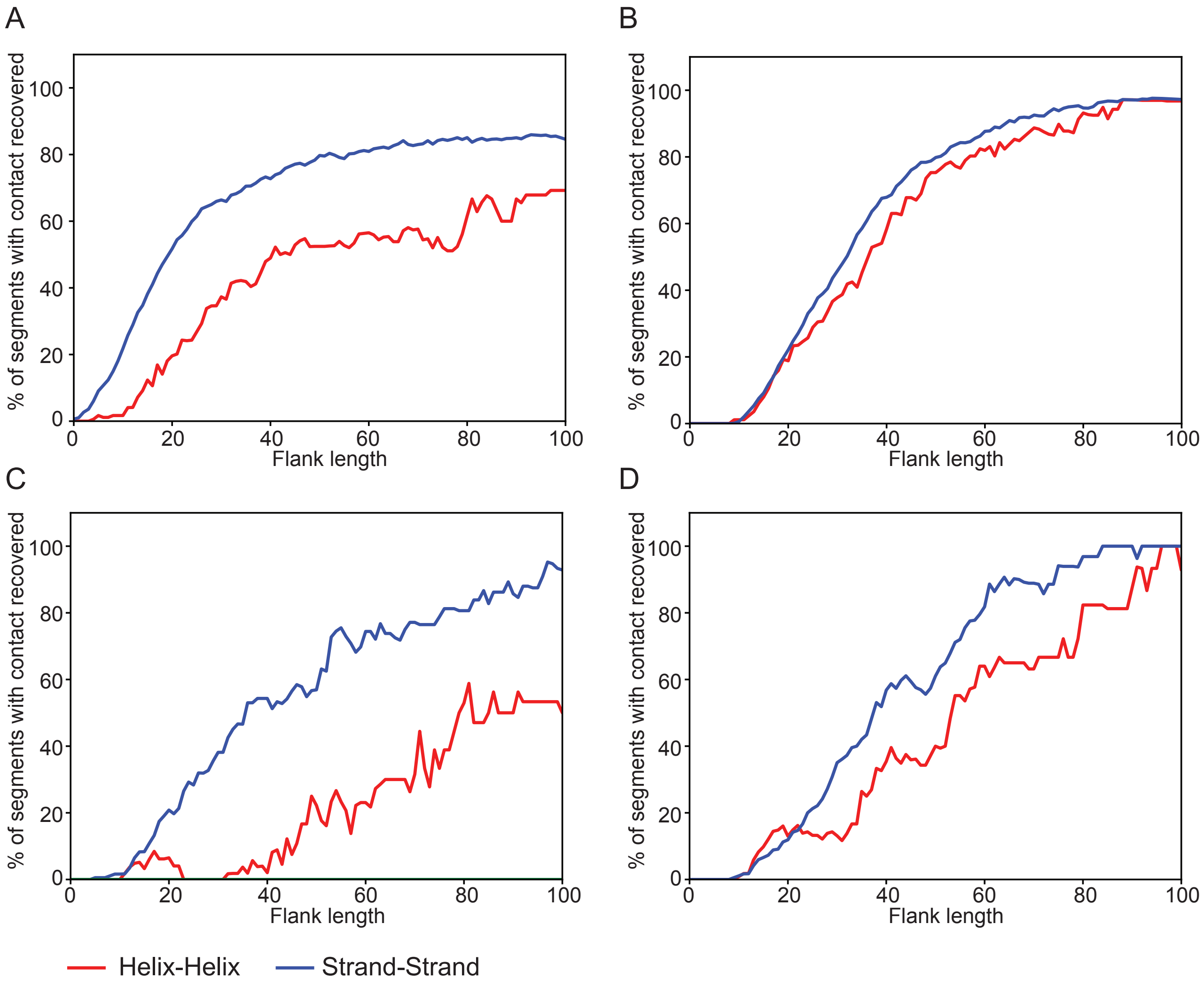


**Fig. S6.** Recovery of strand-strand contacts required less flanking region than helix-helix contacts for contact recovery experiments done with (A) position offset and SSE pairs separated by 50-100 residues, (B) masking and SSE pairs separated by 50-100 residues, (C) position offset and SSE pairs separated by more than 100 residues, (D) masking and SSE pairs separated by more than 100 residues.

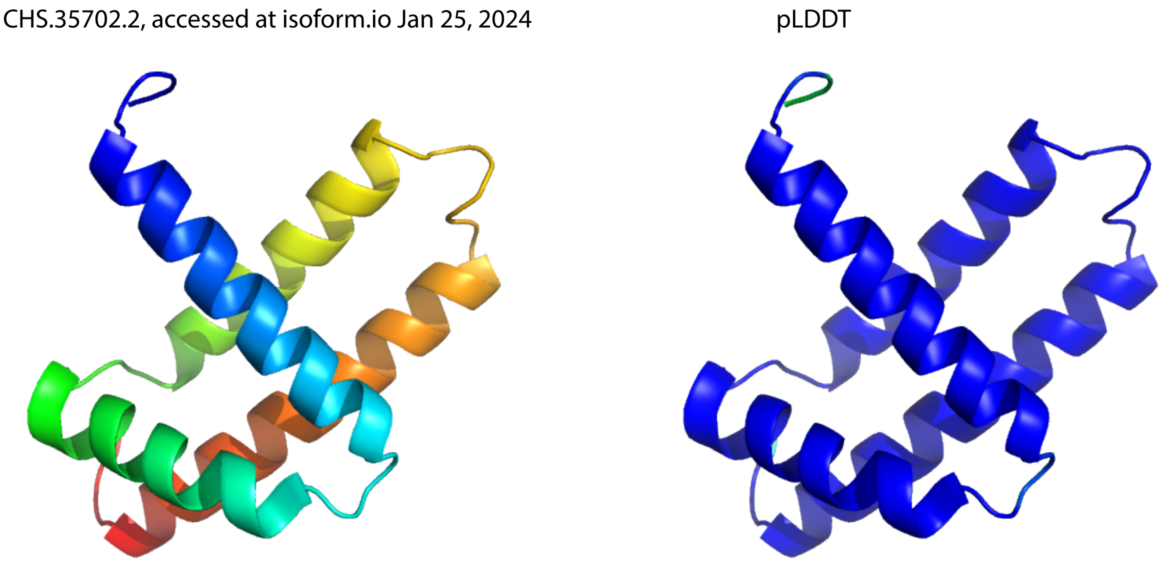


**Fig. S7.** Erroneous prediction for myoglobin isoform CHS.35702.2 is present in isoform.io database, accessed January 2024. (cf. Fig. 2A).

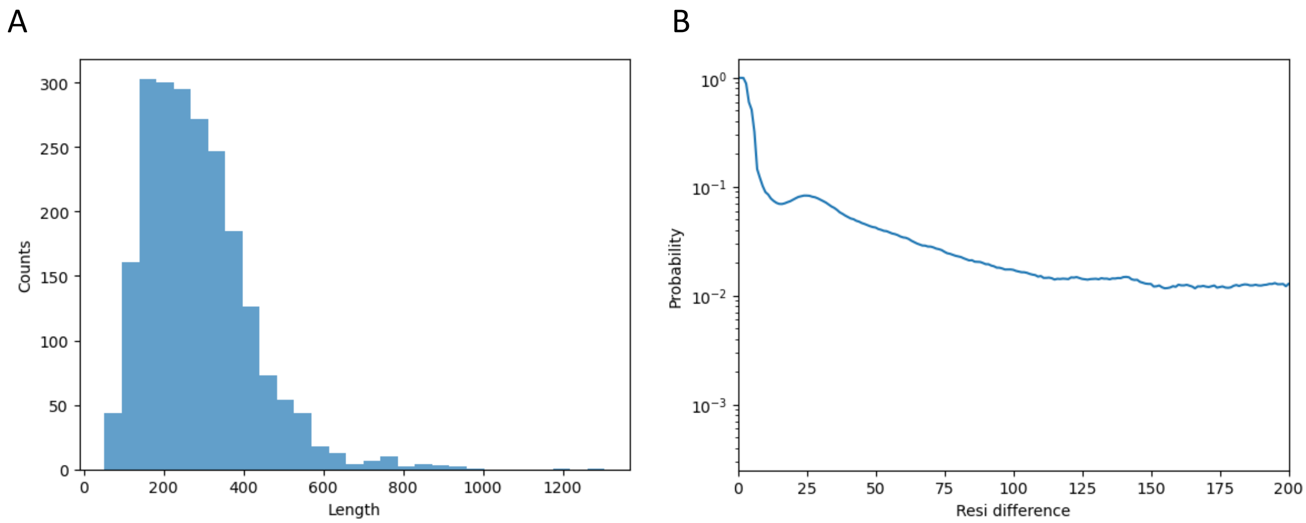


**Fig. S8.** (A) Size distribution of proteins in the dataset. (B) Probability of contact between residues separated by different distances.

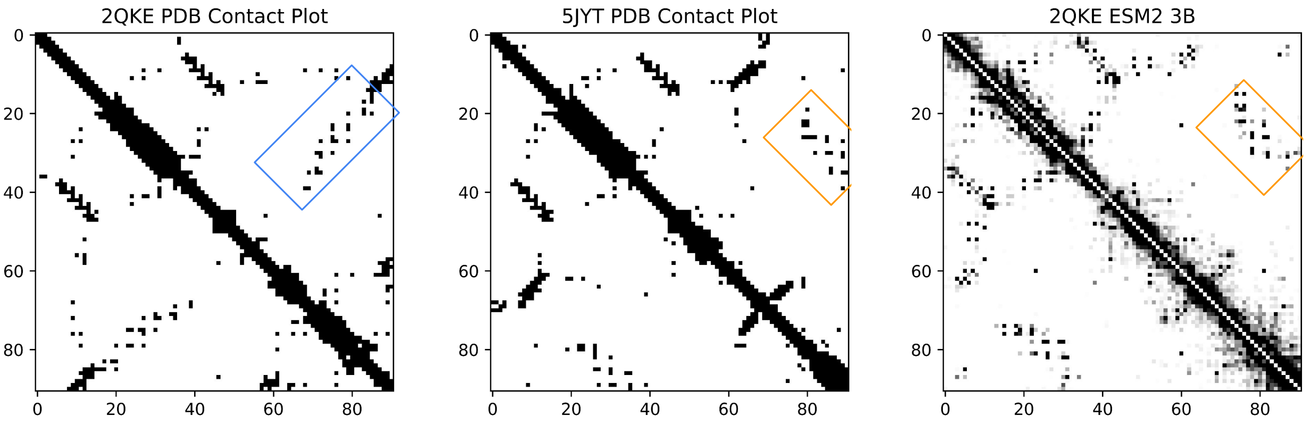


**Fig. S9.** KaiB from *Thermosynechococcus elongatus* thermodynamically favors the Ground state represented in (PDB: 2QKE, distinguishing features boxed in blue), yet also samples the thermodynamically unfavored fold-switched (FS) state (PDB: 5JYT, distinguishing helix-helix interaction boxed in orange). Contact predictions from ESM2 correspond predominantly to the FS state.

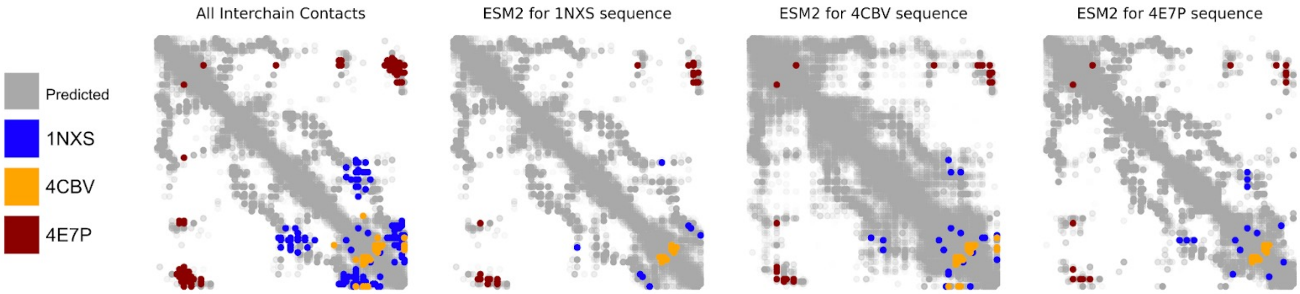


**Fig. S10.** ESM2 predicted contacts compared to the actual interchain contacts of 3 different sequences.

### References

1. HK Wayment-Steele, S Ovchinnikov, L Colwell, D Kern, Prediction of multiple conformational states by combining sequence clustering with AlphaFold2, (Biochemistry), preprint (2022).
2. Malinverni, Barducci, Coevolutionary Analysis of Protein Subfamilies by Sequence Reweighting. *Entropy* **21**, 1127 (2019).
